# GAS Power Calculator: web-based power calculator for genetic association studies

**DOI:** 10.1101/164343

**Authors:** Jennifer Li Johnson, Gonçalo R. Abecasis

**Affiliations:** Department of Biostatistics and Center for Statistical Genetics, University of Michigan, 1420 Washington Heights, Ann Arbor, 48109

## Abstract

**Motivation:** Statistical power calculations are crucial in designing genetic association studies. They help guide tradeoffs between large sample sizes and detailed assessments of genotype and phenotype, help determine which studies are viable, and help interpret research findings. To facilitate widespread use of power analysis in the design and interpretation of genetic studies, it is important to enable users to calculate power and visualize the effect of different models and design choices in convenient, interactive tools that are easily accessible.

**Results:** We developed the Genetic Association Study (GAS) Power Calculator to provide users with a simple interface that can be compute the power of genetic association studies in a convenient browser based interface.

**Availability:** The GAS Power Calculator can be accessed from the web interface at http://csg.sph.umich.edu/abecasis/gas_power_calculator/. Source code is available at <u>https://github.com/jenlij/GAS-power-calculator</u>.

## 1 Introduction

Statistical power is the probability that a study will detect a true effect when there is one – for example, when attempting to establish a connection between a genetic variant and a disease of interest (Purcell *et al*, 2003). Power depends on several factors and calculating it plays a vital role in designing and interpreting scientific studies. In modern genetic studies, power considerations can guide choices between whole genome sequencing (which accesses all genetic variation but is relatively expensive), exome sequencing (which accesses only coding variation and is intermediate in cost) and array genotyping (which accesses mostly common variation but is relatively affordable) (Skol *et al*, 2006; Goodwin *et al*, 2016; McCarthy *et al*, 2008). Calculating the power of a study can help interpret published findings, as interpretation should be different for large, adequately powered studies than for small underpowered studies (Ioannidis, 2005). Recognizing the importance of power in study design and interpretation, granting agencies routinely require power calculations to demonstrate that a proposed study is viable and likely to succeed (Purcell *et al*, 2003). In designing the Genetic Association Study (GAS) Power Calculator, our goal was to make accurate and informative genetic association study power calculations accessible to any scientist. We adapted the widely used algorithms from the CaTS power calculator for two stage association studies (Skol et al. 2006) to work in a modern browser environment and to focus on the types of studies that are now common. The original CaTS tool, which has been used in nearly 1,000 studies (per Google Scholar, as of July 2017), relies on older Windows and Macintosh interactive frameworks that are no longer supported in modern operating systems. Our new GAS implementation works on modern browsers and includes built-in plotting functionality to help users understand the impact of different model parameters and design choices on the power of their study.

## 2 Methods

### 2.1 Features and Functionality

The GAS Power Calculator uses a JavaScript web interface comprised of three sections: inputs, graphs, and results. Users describe the study design by selecting the number of cases and controls, target significance level (which typically depends on the number of markers that will be tested for association), the disease model (multiplicative, additive, dominant, or recessive), prevalence, and allele frequency, as well as genotype relative risk. The calculator then performs relevant computations (detailed on the website) and displays the estimated power, the expected disease allele frequency in cases and controls, the probability of disease for different genotypes, and the frequency of those genotypes. These results are calculated using algorithms adapted from the original CaTS Power Calculator C++ code. The power is computed using implementations of the standard normal based on Hill, (1973) and of the inverse normal distributions based on Wichura (1988). All the source code is freely available on GitHub.

In the graphs section, users can select any of the input parameters to see how its range of values impact power, while the remaining parameters are held constant. This allows, for example, users to graphically explore the consequences of increasing the number of controls, of focusing on rare versus common variants, or of changing significance thresholds. This feature is useful in determining which variables most influence statistical power. The graphs are computed at pre-selected data points that cover the range of the independent variables.

### 2.2 Uses

The following example is a use case for the GAS Power Calculator (Fig. 1). Suppose a user is planning a genome-wide association study with 1,500 cases and 1,500 controls. The plan is to genotype these samples on 300,000 independent SNPs and the user is willing to tolerate a genome-wide false positive rate of 3 (Skol, 2006). Therefore, the target significance level will be 3/300,000, or 0.00001. From previous studies, the user estimates that the disease prevalence is 0.10. Assuming an allele frequency of 0.30 in the general population and a multiplicative model for disease risk, the user wants to determine the genotype relative risk that will result in a power of 80%. To do so, the user enters the fixed parameters and then selects “Genotype Relative Risk” (GRR) as the independent variable to plot against power. Quickly, they learn that a power of 80% requires genotype relative risk of ∼1.30, which they can then judge to be reasonable (or not) for the trait of interest.

**Figure 1.**
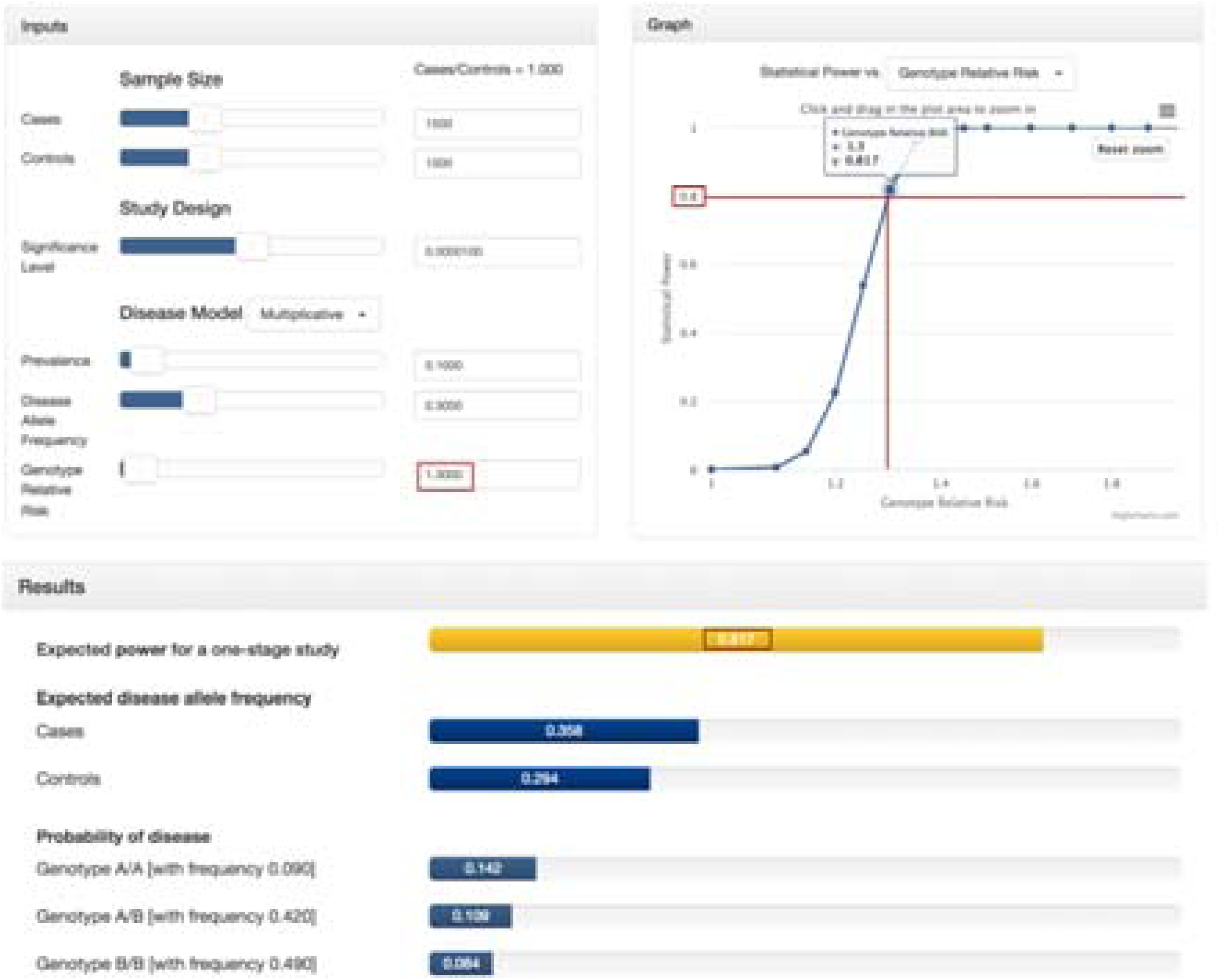
Example screen shot for the GAS power calculator. User specified settings are described in the “Inputs” section in the top-left, graphical summaries can be browsed on the top-right panel, and main power calculation results are summarized in the bottom panel.

Each study will have unique constraints. Typically, we recommend significance thresholds of ∼5x10^-8^ for array-based genomewide association studies (which typically comprise ∼1 million independent tests; McCarthy *et al*, 2008), of ∼5x10^-9^ for sequence-based association studies (which typically comprise additional independent tests; The 1000 Genomes Project, 2015), and of ∼5x10^-6^ for exomewide association studies (which consider only about 1% of the genome; Huyghe *et al*, 2013). Plausible numbers of cases and controls will depend on the resources available to each study and on whether the disease is common or rare. For estimating effect sizes, we recommend users consider the landscape of known genetic findings for other complex traits. Currently, there are many examples of common variants with additive contributions to disease risk and genotype relative risks of ∼1.1 – 2.0, of low frequency variants with modestly higher genotype relative risks of ∼1.5 – 3.0, and of rare variants with even higher genotype relative risks of ∼2.0 – 5.0 (Welter *et al*, 2014).

## 3 Results

We have modernized the original CaTS application to ensure it runs on modern browser environments, converting the original C++ code to JavaScript and incorporating web based interface elements that require no installation. GAS not only performs power calculations, but also allows the user to visualize changes in power over the range of each parameter. The data points used in the plots are hardcoded, therefore the tool currently only has the capability to help the user estimate the power over a range of values, rather than serve as a graphing tool that allows for user uploaded values to plot. We welcome user feedback and suggestions for additional features and improvements. We hope the GAS power calculator will make power calculations for genetic association studies much easier and that it will be useful for many studies to come.

## Acknowledgements

The authors thank Christopher Clark, Kevin Wei Li, and Sean Caron for their invaluable assistance in the web development process. We are also grateful to the Abecasis Lab and our users for their feedback and ideas to improve GAS Power Calculator.

## References

Goodwin S., McPherson J.D., McCombie W.R. (2016). Coming of age: ten years of next-generation sequencing technologies. Nature Reviews Genetics, 17, 333–351. doi:10.1038/nrg.2016.49.

Hill ID. (1973). Algorithm AS 66: The Normal Integral. Applied Statistics, 22, 424. doi:10.2307/2346800.

Huyghe J.R., Jackson A.U., Fogarty M.P., Buchkovich M.L., Stančáková A., Stringham H.M., Sim X., Yang L., Fuchsberger C., Cederberg H., Chines P.S., Teslovich T.M., Romm J.M., Ling H., McMullen I., Ingersoll R, Pugh EW, Doheny KF, Neale BM, Daly MJ, Kuusisto J, Scott LJ, Kang HM, Collins FS, Abecasis G.R., Watanabe R.M., Boehnke M., Laakso M., Mohlke K.L. (2013). Exome array analysis identifies new loci and low-frequency variants influencing insulin processing and secretion. Nature Genetics, 45, 197–201. doi:10.1038/ng.2507.

Ioannidis J.P.A. (2005). Why Most Published Research Findings Are False. PLOS Medicine. doi.org/10.1371/journal.pmed.0020124.

McCarthy M.I., Abecasis G.R., Cardon L.R., Goldstein D.B., Little J., Ioannidis J.P.A., Hirschhorn J.N. (2008). Genome-wide association studies for complex traits: consensus, uncertainty and challenges. Nature Reviews Genetics, 9, 356–369. doi:10.1038/nrg2344.

Purcell S., Cherny S.S., Sham P.C. (2003). Genetic Power Calculator: design of linkage and association genetic mapping studies of complex traits. Bioinformatics 19, 149–150. doi:10.1093/bioinformatics/19.1.149.

Skol A.D. (2006). CaTS Power Calculator. Retrieved March 31, 2017, from http://csg.sph.umich.edu//abecasis/CaTS/.

Skol A.D., Scott L.J., Abecasis G.R., Boehnke M. (2006). Joint analysis is more efficient than replication-based analysis for two-stage genome-wide association studies. Nature Genetics Nat Genet. 38, 209–213. doi:10.1038/ng1706.

The 1000 Genomes Project Consortium. (2015). A global reference for human genetic variation. Nature, 526, 68–74. doi:10.1038/nature15393.

Welter D., MacArthur J., Morales J., Burdett T., Hall P., Junkins H., Klemm A., Flicek P., Manolio T., Hindorff L., Parkinson H. (2014). The NHGRI GWAS Catalog, a curated resource of SNP-trait associations. Nucleic Acids Research, 42, (Database issue): D1001-D1006. doi: 10.1093/nar/gkt1229.

Wichura MJ. (1988). Algorithm AS 241: The Percentage Points of the Normal Distribution. Applied Statistics, 37, 477. doi:10.2307/2347330.

